# Information-based taste maps in insular cortex are shaped by stimulus concentration

**DOI:** 10.1101/774687

**Authors:** E Porcu, KM Benz, F Ball, C Tempelmann, M Hanke, T Noesselt

**Affiliations:** Department of Biological Psychology, Otto-von-Guericke-University, Magdeburg, Germany; Center for Behavioral Brain Sciences, Otto-von-Guericke-University, Magdeburg, Germany; Department of Neurology, Otto-von-Guericke-University, Magdeburg, Germany; Institute of Neuroscience and Medicine, Brain & Behaviour (INM-7), Research Centre Jülich, Jülich, Germany; Institute of Systems Neuroscience, Medical Faculty, Heinrich Heine University Düsseldorf, Düsseldorf, Germany

**Keywords:** gustatory, human, fMRI, MVPA, concentration

## Abstract

Taste processing is an essential ability in all animals signaling potential harm or benefit of ingestive behavior. Although the peripheral taste coding is well understood, current evidence for central taste processing remains contradictory. To address this issue, human participants judged pleasantness and intensity of low and high-concentration tastes (salty, sweet, sour, bitter) in two fMRI-experiments. High-resolution fMRI and multivariate pattern analysis were used to characterize taste-related informational content in human gustatory cortex (GC). Clusters within GC were narrowly tuned to specific tastants consistently across tasks. Importantly, taste concentrations completely altered the spatial layout of putative taste-specific maps with distinct, non-overlapping patterns for each taste category at different concentration levels. Together, our results point at population-level representations in human GC as a complex function of taste category and concentration.

## Introduction

In mammals, taste identification constitutes a critical ability to ensure survival through selection of nutritional versus potentially harmful food. Although humans have developed a fine ability to distinguish different tastes and taste compounds, it is yet unclear how basic and complex tastes are distinctively represented in the gustatory cortex (GC).

Other sensory cortices are traditionally characterized by specific topological organizations of neurons arranged into specialized functional maps, e.g. orientation maps in the primary visual cortex or tonotopic maps in the primary auditory cortex. Hence, one might hypothesize that there is a functional map type of organization (Wilson and Bednar, 2015) in chemosensory cortices as well, based on basic taste categories. Yet, it still needs to be proven whether any general topological principle applies to gustatory cortices (see Giessel and Datta, 2014 for olfaction).

Only recently have putative gustotopic maps been observed in rodents (e.g. Chen et al., 2011). There, pools of neurons were characterized by direct associations with specific taste qualities (salty, sweet, bitter and umami) within specialized sub-regions of mouse’s insular cortex. In clear contradistinction however, the majority of animal studies reported overlapping clusters of neurons broadly tuned to distinct tastes (e.g. Accolla et al., 2007). For instance, Fletcher and colleagues (Fletcher et al., 2017) showed that neurons were preferentially tuned to specific tastes or combination of tastes; nonetheless such neurons were intermingled with neurons tuned to different tastes, in line with the notion of a non-topological gustatory organization in other species (for similar results in non-human primates, see Scott and Plata-Salamán, 1999).

In humans, to date, the organization of the GC remains poorly understood. The putative human GC, located in anterior to middle insula (Iannilli et al., 2014; Veldhuizen et al., 2011), is inherently a multisensory area which exhibits a wide range of responses not exclusively related to gustatory stimuli (e.g. somatosensory, thermal responses). This organization makes particularly challenging to isolate activity elicited by taste stimuli from other influences, e.g. somatosensory responses.

In accord with the vast majority of animal data, most human imaging studies have reported a scattered organization of neuronal populations broadly tuned to different taste categories, with no actual evidence of dedicated taste-specific maps. Only few fMRI-studies (Prinster et al., 2017; Schoenfeld et al., 2004) have reported taste-specific clusters of voxels which may resemble a gustotopic type of organization based on mass-univariate analyses. However, both studies were penalized by small sample sizes, passive tasting (i.e. lack of task) and presentation of only one intensity per taste. The latter two points strongly limit the generalizability of these findings as intensity variations can dramatically alter population responses to tastes (Canna et al., 2019; Small et al.,2003) as can the task at hand (Grabenhorst et al., 2008). Moreover, studies exclusively relying on mass-univariate analyses suffer from the common inability to discriminate *spatial patterns predicting* specific taste categories, tasks and intensities and thus the ability to assess the informational content within activated clusters. To overcome this impasse, Chikazoe and colleagues (2019) have recently adopted a multivariate approach to classifying basic taste categories as well as different chemical compounds of the same taste categories. While these authors were able to identify common patterns of activities in the insular cortex associated with basic taste qualities, irrespective of chemical compounds, the lack of variation in tastant concentration and task compromises the conclusion that a taste-specific map indeed exists. Moreover, the absence of an adequate measure to quantify the selectivity of specific patterns for particular taste categories further impairs their interpretation. Hence, the aim of the present study was to test for a general *gusto-topic* organization and, more specifically, to provide a comprehensive characterization of the human gustatory cortex as a function of (I) taste quality (bitter, salty, sweet, sour), (II) taste concentration (high, low) and (III) task (pleasantness, intensity judgements). To this end, 24 participants were tested in two fMRI experiments in which we administered four basic taste categories at two different concentrations plus an additional neutral compound. Participants judged intensity in one experiment and pleasantness in the other experiment. Crucially, a spatially constrained classifier (searchlight approach) was trained on data of the first experiment and tested on the second experiment and vice versa, to identify common, task-independent yet taste-specific representations. Moreover, taste-specific tuning functions were calculated for each searchlight to address the issue of narrow vs. wide taste-specific tuning functions in human GC. Finally, across- and within-concentration classification analyses were used to assess the influence of concentration on taste-specific representations.

## Results

### Behavior

Participants were able to correctly perceive taste identity in the majority of all cases (see supporting Fig. S1a-b). Moreover, they rated taste valence differently depending on taste concentration and taste identity, as testified by the significant interaction (*F*(4, 23) = 3.028, *p* = 0.0186; η^2^ = 0.030): with sweet always showing significantly enhanced and bitter showing significantly decreased ratings compared to neutral (see supporting Tab. S1 and supporting Fig. S2c for details). Concerning the intensity judgments, participants perceived a consistent difference in intensity between high and low concentrations, as expected (main effect of intensity: *F*(1, 23) = 34.398, *p* < .001; η^2^ = 0.181). Moreover, a main effect of taste identity was observed (*F*(4, 23) = 9.810, *p* < .001; η^2^ = 0.280), while the interaction between intensity and taste identity did not reach significance (*F*(4, 23) = 2.237, *p* > .05; η^2^ = 0.048, see supporting Tab. S1 and supporting Fig. S1d for details).

### Neural underpinnings

The central focus of this study was to characterize the functional organization of putative GC, thus left and right insular cortex were targeted by means of ROIs derived from independent functional parcellation of the insula (Fan et al., 2016). Sparse multinomial logistic regression (Krishnapuram et al., 2005) was used to train and test in turn on the first experiment and second experiment through a searchlight approach (Kriegeskorte et al., 2006).

### Cross experiment decoding

First, we isolated fMRI patterns associated with taste categories. Given previous reports on the influence of taste concentration on fMRI signals (e.g. Canna et al., 2019), separate classifications were calculated for high and low intensities. These critical comparisons will unveil task-invariant yet single or multiple taste-preferring patterns if present.

#### High concentration

Classification among high concentration tastes revealed patterns of spheres for all four taste categories which were successfully decoded across the two experiments (see Fig. 1a, right side). For this analysis, we first estimated probabilities (rather than accuracies) to quantify the selectivity of each searchlight sphere for a given taste quality. This set of probabilities provides a continuous multidimensional measure describing the selectivity of each searchlight sphere (one probability for each taste) rather than a “winner-take-all” accuracy measure (Anzellotti and Coutanche, 2018). After initial classification, we then estimated evidence ratios (ER) by contrasting correct classifications with false positives (i.e. higher probability of a particular taste when another taste was presented, see Fig. 1a for significant ER maps, p(cluster) < .05, FWE-corrected, with an auxiliary threshold of p(voxel) < .001). Statistical maps of ER were subsequently used to assess the selectivity of each sphere, based on a tuning index. In particular, tuning indices (range 0-1) were computed based on the evidence ratio for a particular taste relative to the highest evidence for all other tastes. Indices equal or larger than 0.5 indicate an increasingly narrow tuning for a specific taste category (see Methods section for further details). Remarkably, our results provide evidence for a consistent narrow selectivity for all taste categories across experiments, i.e. independent of task (see Fig. 1b for tuning maps).

**Figure 1.**
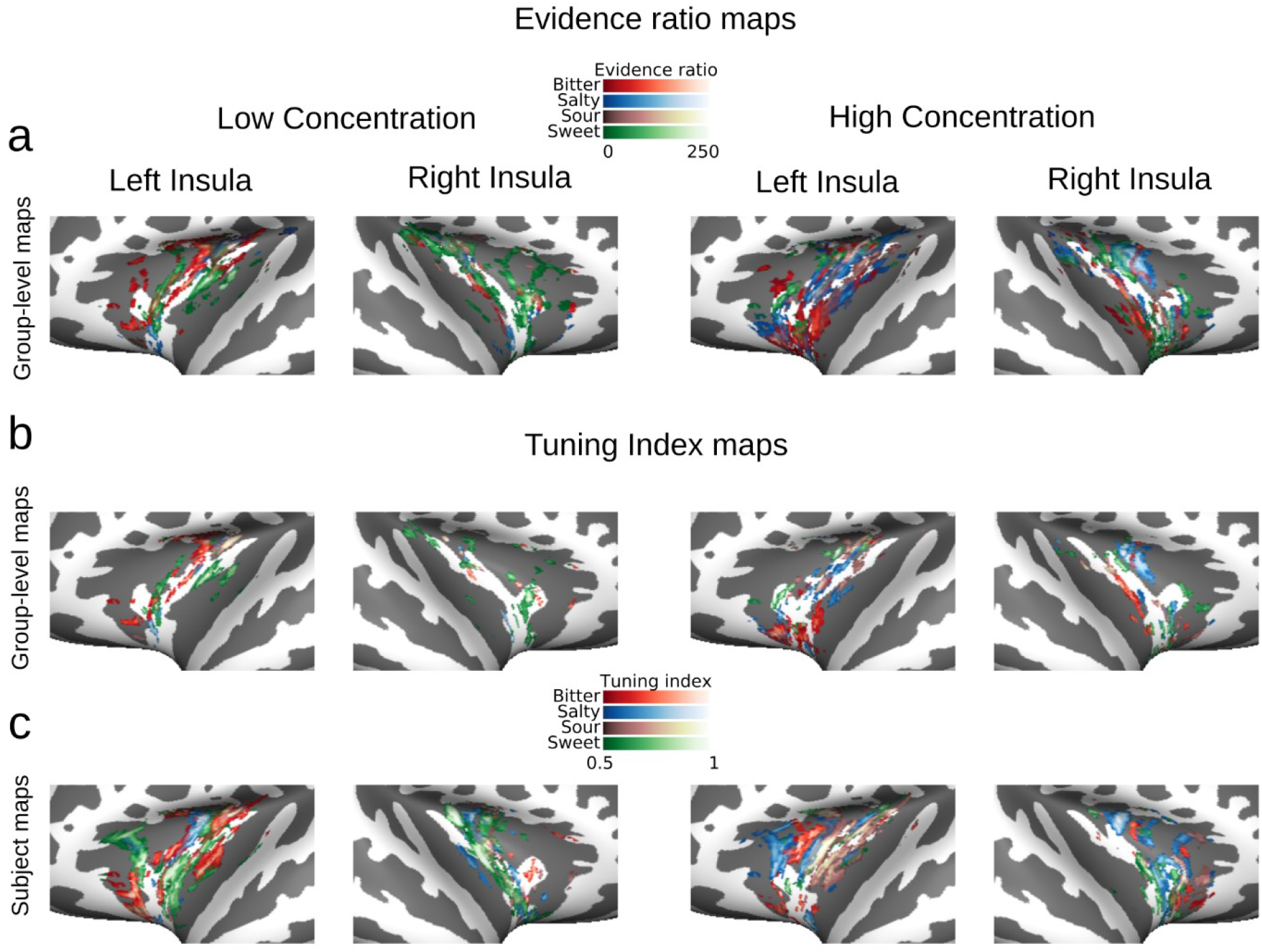
Cross experiment taste maps: (a) Flat maps depict significant clusters of evidence ratios (cluster-thresholded at p_FWE_ < 0.05 with an auxiliary voxel-threshold of p < 0.001) in insular cortex at the group-level, obtained through cross experiment decoding. Plots were created by projecting 3D brain voxels onto a 2D surface via Nilearn 0.5.0 (Abraham et al., 2014)and subsequently plotting the brain surfaces by using Visbrain 0.4.0 (Combrisson et al., 2017). Note that ERs for particular tastes can and do overlap. (b) Tuning maps depict spheres narrowly tuned to single tastes at the group level (thresholded at > 0.5). (c) Tuning maps for an illustrative subject. The information provided by group level maps may be misleading given the extreme variability of the insula’s functional microstructure (Prinster et al., 2017; Schoenfeld et al., 2004). Single subject maps exhibit a stronger degree of spatial continuity for specific taste categories than the group-level maps. Left columns always represent low concentration tastes, right columns represent high concentration tastes.

Moreover, the number of spheres narrowly tuned to one particular taste varied, with bitter showing consistent effects in ~4.3 % of insular spheres, salty in ~3.5 % of insula’s spheres, while sweet and sour registered ~2.8 % and ~1.8 %, respectively. Similar sphere counts were obtained in both left and right insula. Next, we tested for spheres broadly tuned to more than one taste. However, virtually none of the clusters coding a combination of two or three tastes survived the selection criteria.

To confirm at the single-subject level that these sphere-based observations are not simply due to large interindividual differences in topologies due to group-mean voxelbased averaging (Prinster et al., 2017; Schoenfeld et al., 2004), we next tested for differential effects of taste category based on single subject results (see Fig. 1c for tuning maps of an illustrative subject). Here, 8.89% (median +/− 1.2 % standard error), 8.40% +/− 1.2 %, 8.38% +/− 1.2 %, 6.75% +/− 1.1% of insular spheres averaged over hemispheres responded to salty, bitter, sweet and sour, respectively (see Fig. 2a, leftmost columns for all scatter plots of all individual subjects).

**Figure 2.**
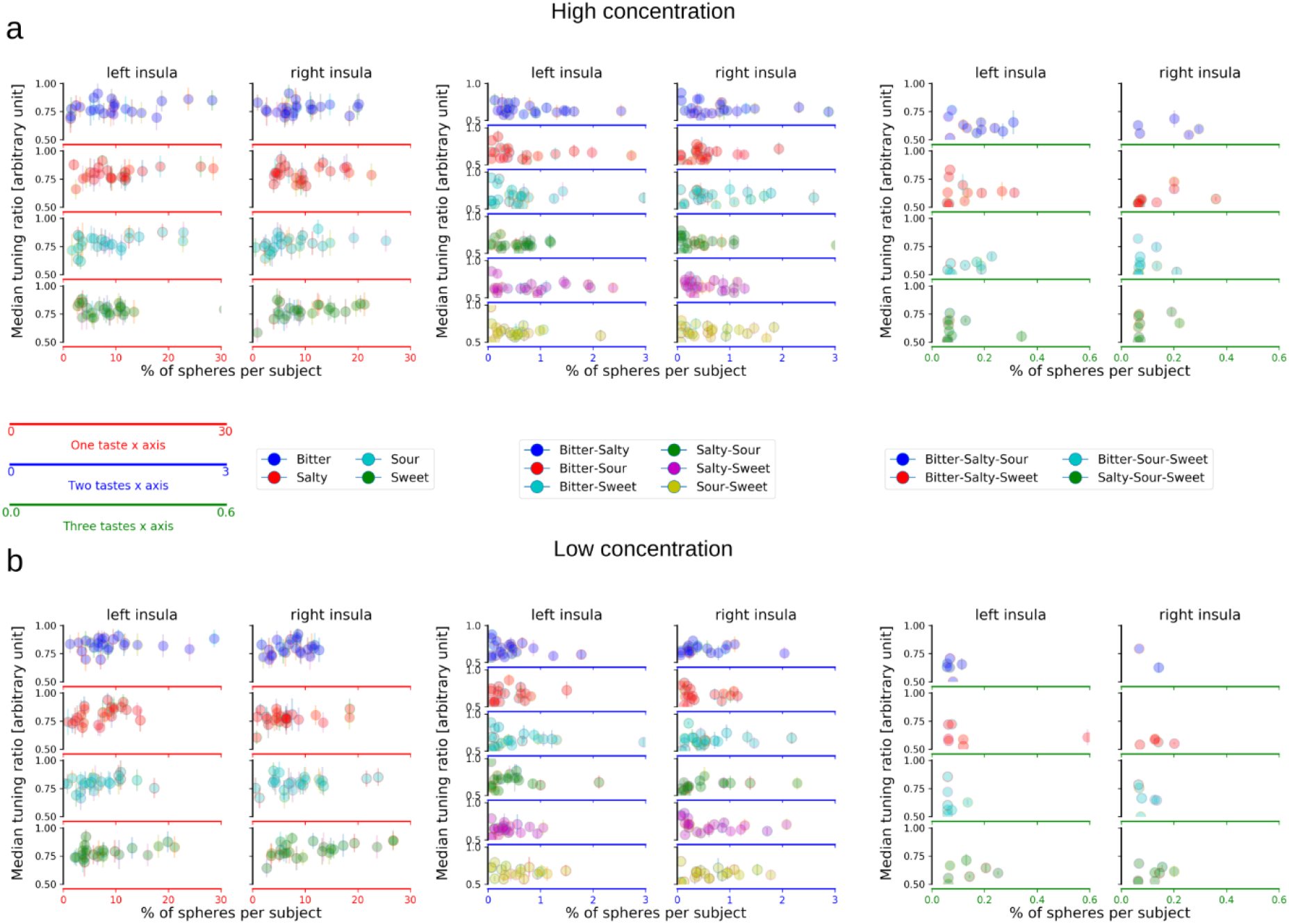
Tuning indices at subject level: Scatter plots depict median tuning index for all subjects separately for high (a) and low (b) concentrations. The y-axis represents the median across tuning indices for a specific taste or a compound of tastes. The x-axis indicates the percentage of spheres that shows a preferential tuning for a taste or a compound of tastes. Scatter plots, from left to right, represent percentage of spheres tuned to single tastes (2 leftmost columns), double tastes (2 center columns) and triple tastes (2 rightmost columns); the left and right columns of each taste configuration represent the left and right insula, respectively. Note that the x-axis is scaled differently for single-, double- and triple taste spheres (leftmost, middle and rightmost columns) due to the decreasing number of spheres responsive to double and triple taste compounds.

All subjects expressed taste-selective spheres for all tastants. No significant differences in sphere count were observed for hemisphere, taste category or their interaction (p’s > .4 and higher) mirroring the group-level results (though note that subject-specific sphere counts are higher than group-level counts, indicating increased sensitivity). Moreover, the number of spheres was not related to perceived intensity or pleasantness for any taste (p >= .424, Bonferroni-corrected). Finally, lower counts were found for two-taste (0.5% +/− 0.3%) and three-taste preferring spheres (see Fig. 2a, middle and rightmost columns; more than half of the subjects did not show any significant three-taste preferring clusters; see also supporting Tab. S2a-b). No significant differences across hemispheres or taste category were observed for the double or triple-tuned spheres, confirming the pattern observed at the group level.

#### Low concentration

As second critical comparison, we tested for task-invariant taste-specific patterns at low concentration. Figure 1a (left side) depicts significant clusters of evidence ratio maps for each taste category for the low concentration stimuli. As for high concentration maps, we again observed task-invariant yet taste-selective clusters for all four tastes (see Fig. 1b). Overall cluster sizes were comparable to those observed for the high concentration conditions at the group level. In particular, the subjects-specific sphere count for the low-concentration conditions revealed an increase in insular spheres tuned to sweet (~4.7 %), while bitter (~2.3 %), salty (~0.7 %) and sour (~1.5 %) preferences slightly decreased relative to high concentration spheres. As for the high concentration, virtually none of the clusters were tuned to more than one taste category. Again, confirmatory analyses of single-subject results corroborated the group level results (see Fig. 2b), with higher counts for single taste preferring spheres (bitter: 8.27% +/−1.0; salty: 6.88% +/−.9; sour: 7.47% +/− 1.0; sweet: 9.68% +/− 1.3) than two- and three-taste preferring spheres (see supporting Tab. S2). Moreover, subjective intensity or pleasantness ratings were again not related to subject-specific sphere counts for any taste category (p >= .512, Bonferroni corrected). Together, the findings for both concentration levels reveal that there are consistent taste-preference clusters in the insula and that most taste-sensitive spheres expressed a preferential tuning to one taste only. However, these analyses above were independent from each other, thus cannot answer the critical question whether the observed task-independent taste-specific subregions were also independent of taste concentration.

### High versus low concentrations

The most critical comparison therefore tested whether the observed task-independent clusters were also not affected by stimulus concentration. For this direct fine-grained analysis of the relationship between high and low concentration maps, we quantified the amount of overlap between high and low concentration maps for both group and single subjects maps for all taste categories. Specifically, we counted the number of spheres coactive in low and high concentration tuning maps and we then computed the ratio between the number of coactive spheres using geometric mean between the two compared cluster sizes. Geometric averaging was adopted to account for the overall different size of the clusters of interest.

Otherwise, the chance to observe high or low overlap might have been biased by the relative size of each cluster (e.g. two bigger clusters would have had a higher probability to share common spheres than two small clusters). This procedure provided a continuous index of overlap ranging between 0 and 1, where 0 indicates no overlap and 1 indicates complete overlap. We computed the index of overlap separately for each taste to determine to which extent spheres that showed a preference for a specific taste at a low concentration showed a preference for the same or a different taste at a high concentration. Remarkably, the overlap between high and low concentration maps - obtained from the cross experiment classification - were extremely low suggesting distinct taste-related patterns as a function of stimulus intensity (see Fig. 3a middle diagonal, bitter left insula: 0, bitter right insula: 0, salty left insula: 0.03, salty right insula: 0, sour left insula: 0.05, sour right insula: 0, sweet left insula: 0, sweet right insula: 0.04, see supporting Fig. S2 for virtually identical results at the single-subject level). Remarkably, some spheres even switched their taste-specific preference (e.g. from bitter to sour as indicated by the off-diagonal cells).

**Figure 3.**
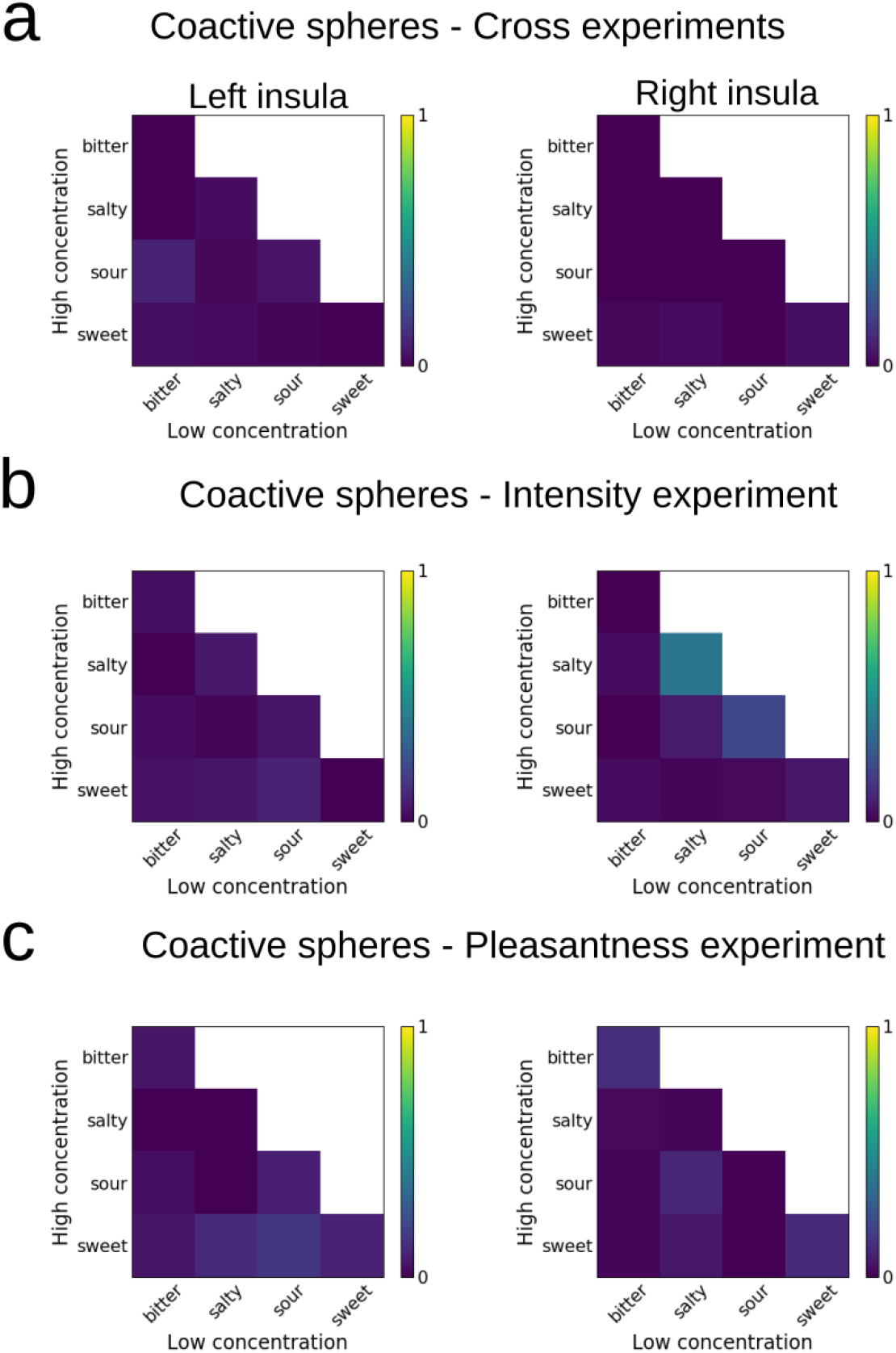
Coactive spheres across taste concentrations: (a) Heat maps depict the coactivity index of spheres based on data from narrowly tuned spheres for the cross experiment decoding. Coactivity is expressed as an index of overlapping spheres across concentrations, where 0 indicates the absence of overlap and 1 complete overlap. (b) Heat maps represent the coactivity index based on classifications for intensity judgements. (c) Heat maps represent the coactivity index for pleasantness judgements. The main diagonal represents coactive spheres coding for the identical taste category across concentrations. Values outside the main diagonal indicate a switch in taste preference as a function of concentration.

To test the exact response properties of the spheres that exhibited concentration-invariant taste-specificity, we visualized the amount of agreement between high and low-intensity sphere values using Bland and Altman plots (Bland and Altman, 1999). Specifically, the plots in Figure 4 show the difference between sphere values for high and low concentrations (see the Methods section for further details). Here, deviations from zero on the y-axis indicate the amount of disagreement between high and low intensity sphere values, the closer to zero the less sphere values disagree. There was only a small subset of coactive spheres between high and low concentrations with 4-8 participants per taste showing no concentration-invariant clusters at all (though note that these concentration-invariant spheres in the responsive participants shared particular properties as they were grouped together in the Bland-Altman-plots along the y-axis).

**Figure 4.**
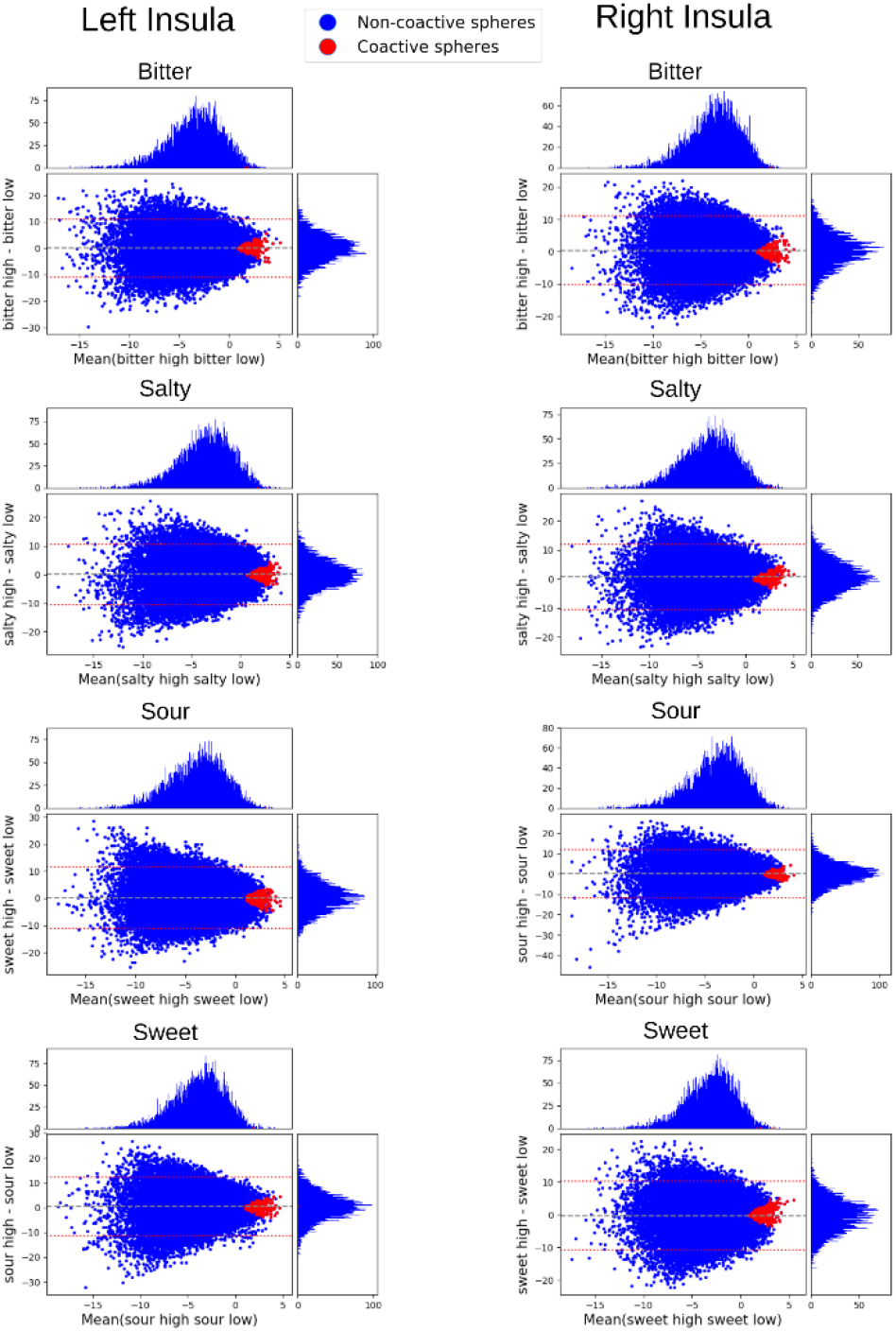
Coactive concentration-independent spheres: Bland and Altman plots represent the amount of agreement between low and high concentration spheres for the cross experiment classification. Blue dots represent the agreement of high and low concentration spheres in the insula for all subjects. Red dots represent the amount of agreement for coactive spheres of all subjects. The clustering around zero (y-axis) suggests non-trivial similar evidence ratios in low and high concentrations, indicating concentration independent spheres. The x-axis indicates the average between high and low concentration of log transformed evidence ratios. The y-axis indicates the difference between high and low concentration log transformed evidence ratios (see Bland and Altman plots section for further details). The top-distribution represents the mean between high and low concentrations for a given taste. The left-distribution represents the difference between high and low concentrations for a given taste.

Finally, we tested whether the few concentration-invariant spheres anatomically group together in a particular region of the insula. Since the insula contains many higher-order gustatory and multisensory regions in addition to the primary gustatory cortex (PGC), we reasoned that concentration-invariant spheres might be clustered in PGC. To test this, we first generated group mean maps for each taste. However, no location of insula contained concentration-invariant spheres from more than three participants. Second, we checked whether any particular subfield within the insula contained most of the subject-specific concentration-invariant spheres using the subfield definition provided by Fan and colleagues (Fan et al., 2016). However, concentration-invariant spheres were present in all subfields but were usually found in less than half of all participants per subfield (see supporting Fig. S3). Thus, we observed no anatomical hotspot for concentration-invariant spheres.

### Within experiments decoding

In order to further confirm that the topological distinction observed between low and high concentrations was consistent in each single experiment and not merely due to task differences across the two experiments, we performed four additional classifications, i.e. we separately decoded high and low concentrations within each experiment. For this analysis, the cross-validation was based on a different training and test set: a leave-one-out procedure was used in which the classifier was trained on three out of the four runs and tested on the remaining one.

As expected, we were again able to decode patterns of spheres, here within the individual experiments, which were associated with each taste at high and low concentrations (see Fig. 5a-b). In Figure 3b and 3c, we show the index of overlap between high and low concentrations separately for each experiment; as for the cross-experiments maps (see Fig. 3a), we see very little overlap between high and low concentration maps in the within-experiment analyses.

**Figure 5.**
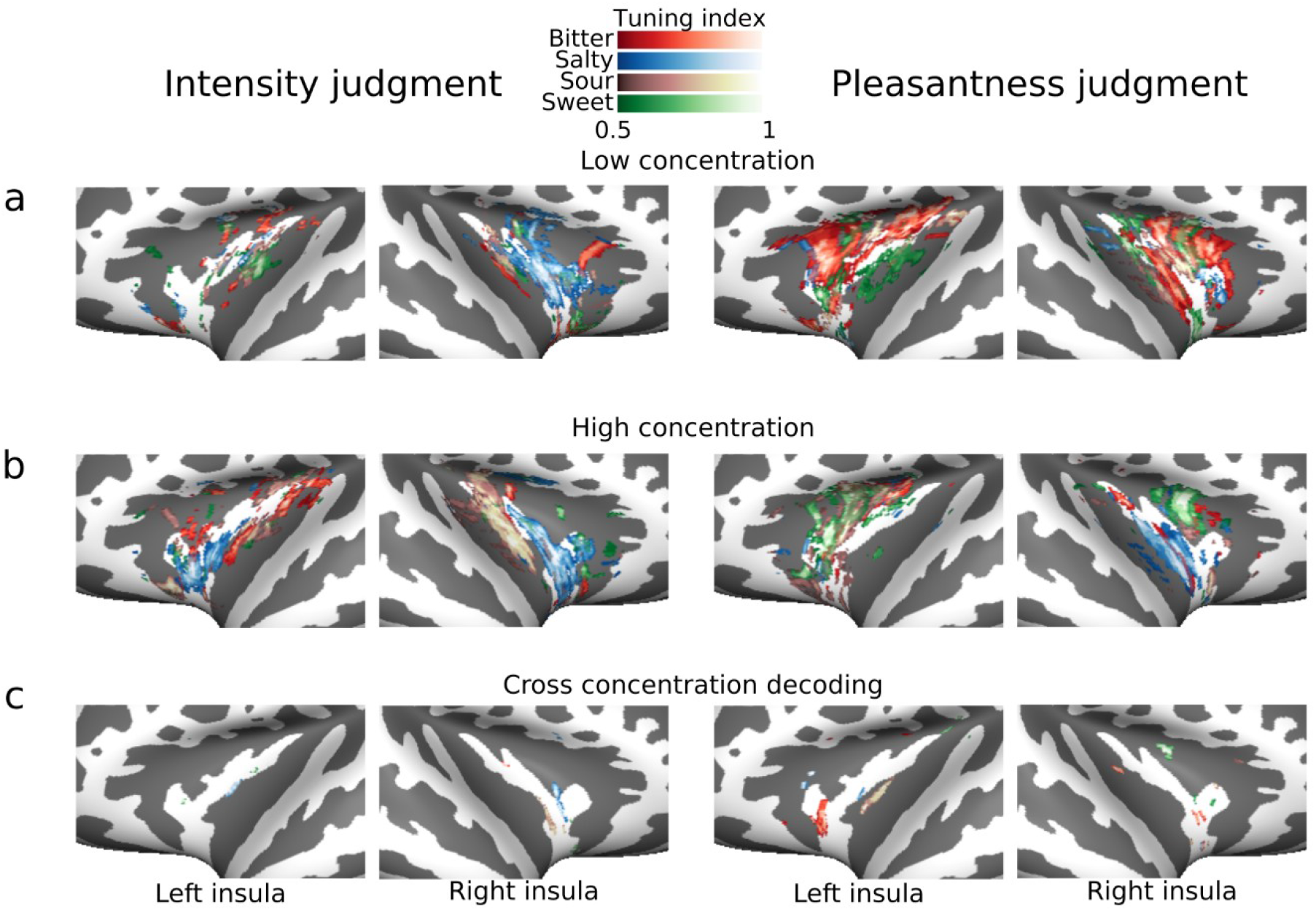
Within experiments tuning maps: (a) Left and right flat maps depict narrowly tuned spheres in insular cortex for low concentration tastes, separately for intensity judgements (left) and pleasantness judgements (right). (b) Left and right maps depict narrowly tuned spheres in insular cortex for high concentration stimuli, for intensity and pleasantness judgements, respectively. (c) Left and right plots depict maps representing tuning indices for cross concentrations decoding for each taste category, separately for intensity and pleasantness judgements. Note that within concentration decoding produces consistent results (a-b), whereas cross-concentration decoding (c) reveals almost no consistent clustering.

### Cross-concentrations decoding

Lastly, in order to confirm whether high and low concentration maps reflect truly distinct populations, we performed two additional classifications, one per experiment. We trained the classifier on the low concentration tastes and tested on the high concentration tastes and vice versa for each experimental session.

We reasoned that if different populations are coding for low and high concentrations, we should observe a poor classification. In line with our hypothesis and the previous decoding results, we indeed observed only a small amount of spheres coding indiscriminately for low or high concentration tastes (Fig. 5c): Based on the tuning maps, in experiment 1, bitter exhibits only 0.1% of narrowly tuned spheres, salty: 0.5%, sour: 0.15% and sweet: 0.17%; similar results are obtained in the experiment 2, where bitter exhibits 1.2%, salty: 0.15%, sour 0.5% and sweet: 0.4%. Together, this pattern of results is in clear disagreement with the notion of broad concentration-invariant maps of taste qualities in the human insula.

## Discussion

The main objective of the present study was to test for task- and concentration-invariant maps in human gustatory cortex through an information based approach. We tested identical participants in two distinct fMRI experiments on different days and successfully decoded patterns of spheres associated with each specific taste category in all participants. Remarkably, we were able to decode patterns of spheres associated with the specific taste categories across distinct behavioral tasks and time, suggesting robust task-invariant taste representations in all individual participants. Most importantly, low- and high-concentration taste-selective representations showed very little overlap, contradicting the notion of an exclusively topological organization based on linguistic taste labels.

Significantly extending previous observations (Chikazoe et al., 2019), we observed patterns of spheres narrowly, but not exclusively, tuned to specific tastes as well as smaller patterns of more broadly tuned spheres in the left and right insula at the group-level and confirmed taste-specific patterns when considering only one concentration. The information provided by group level maps may nonetheless misrepresent the actual organization observable at the single subject level given the extreme variability of the insula’s functional microstructure as previously pointed out (Prinster et al., 2017; Schoenfeld et al., 2004). When inspecting single subjects’ maps, we observed patterns exhibiting a stronger degree of spatial continuity for specific taste categories. It has to be noted though, that such subject-specific tuning maps showed a mixture of spheres narrowly tuned towards one taste category and others exhibiting a broader tuning while keeping a distinct preference for a specific taste.

As a consequence of the structural variability across individuals, spheres equally tuned to two or three tastes were virtually absent at the group-level. Also at the single subject level, the number of spheres coding two tastes was much smaller than spheres coding single tastes; likewise, only a few subjects showed spheres coding for three different tastes. This lack of patterns equally tuned to different tastes might be ascribed to the stringent criteria that we adopted to identify tuning indices. Alternatively, responses in multi-taste spheres might be more easily elicited by presenting compound tastes which was not part of our current experimental design. In line with our results however, Fletcher and colleagues (Fletcher et al., 2017) also observed single taste preferences in the majority of neurons.

Considering topological theories, we do not find any evidence for common maps across concentrations, though stable clusters were observed for single concentrations in line with previous research (Chen et al., 2011; Prinster et al., 2017). Moreover, we observed a patchy organization in insular cortex for different taste categories rather than a highly ordered topography (Chen et al., 2011; Prinster et al., 2017). Thus, our data rather supports a model of distributed populations of neurons coding for different taste qualities, than a model of highly specialized maps. A population coding model seems to be supported by the distinct topological organization that we observed in high and in low concentration maps (note that even for the low-concentration tastes all behavioral responses differed from each other, pointing to a sufficiently high concentration for perception, in accord with the distinct patterns of neural activity). Especially at the single subject level (see Figure S2 in Supporting material), subsets of the spheres that preferentially coded for a specific taste at low concentration changed their preference to a different taste at high concentration, hence suggesting that taste coding might dynamically change according to relevant features and might be achieved through a distributed population rather than rely exclusively on rigid functional maps. One might argue that the difference in representation of low vs. high-concentration tastes might be due to the fact that participants were less able to identify the low-concentration tastes (relative to the higher ones). However, a confusion between different low-concentration tastes should lead to less stable decoding patterns rather than the stable patterns for each low-concentration taste observed here. Moreover, the lack of overlap for low and high concentration patterns also suggests that the coding mechanisms in the gustatory cortex differs from other, topologically-ordered systems, e.g. vision: There, a change in stimulus intensity may modulate the firing rate of identical neurons and classification accuracy (due to changes in SNR) yet should yield similar overall patterns; whereas with our gustatory results, tuning patterns stay robust (see Fig.2) but seem to change location. In accord, a recent study conducted in mice (Wu et al., 2015) targeted sensory neurons in the geniculate ganglion, hence focused on an earlier stage of taste processing, and found a robust change in preferential tuning when taste concentration was systematically varied. Up to 51% of the narrowly tuned neurons exhibited a taste-selectivity change from narrow to broad tuning, suggesting that at least at an intermediate processing stage mice brains rely on a pattern coding mechanism. Although, we observed a similar scenario at a higher stage of processing, our results seem to provide further evidence in favor of a population coding model.

Finally, the amount of overlapping spheres between high and low concentration maps was consistently low. In some cases, there were no overlapping spheres, thus suggesting that, at least partially, two different underlying populations of neurons might be preferentially coding for a taste category at high and low concentrations. Our observations are also supported by a recent study in rats (Fonseca et al., 2018), which shows that over a large number of neurons only a “small” subset is able to code for both high and low concentrations of sucrose. In accord, a monkey study (Scott et al., 1991), observed that less than 1.5 % of the recorded neurons in the insular cortex exhibited a linear response as a function of the increase in glucose concentration. Although those studies report effects on the cellular level and might thus be difficult to directly link to population-based fMRI results, it has nevertheless been shown for the visual modality that orientation tuning can be picked up with fMRI (Kamitani and Tong, 2005), even though the spatial resolution of fMRI was much coarser than the size of orientation columns. Potentially, a similar effect was observed here.

While only a small number of concentration-independent taste-preferring spheres were found in our study, concentration-dependent taste-preferring spheres were more common and found in all participants. However, the exact spatial layout differed between participants and did not seem to follow a highly organized topological arrangement. It has also been proposed that chemical senses have evolved to detect and evaluate potential primary reinforcers (e.g. Prinster et al., 2017). Thus, the organizational principle in gustatory cortex should be based on valence rather than physical stimulus properties. However, the largest taste-specific effects found here were task-independent and did not vary with perceived pleasantness (or intensity) in contrast to this hypothesis. Hence, we propose that the variation in neural signaling which was linked to valence in previous studies (Grabenhorst et al., 2008; Prinster et al., 2017) may rather be based on changes of a physical property, i.e. concentration, which governs perceived pleasantness and significantly shapes informational content in low-level gustatory cortex.

In summary, our results demonstrate clusters in the human insular cortex preferentially tuned to specific taste-categories in all single subjects. Importantly, variations in stimulus concentration reveal a complex pattern of activity with little overlap between concentration-dependent taste-specific representations. Together, these results point at a population-based organization in gustatory cortex and highlight that gustatory cortex might be coding a complex mixture of taste identity and concentration rather than a representation based solely on taste identity.

## Methods

### Participants

25 healthy volunteers took part in two experiments; one participant was excluded from the final analysis due to a technical problem with taste delivery, hence resulting in 24 participants (16 female, age: M = 24.9, SD = 3.6). Participants fulfilled the following inclusion criteria: no known allergies/sensitivity to the chemical solutions used in the experiments, no respiratory tract infections in the two weeks prior to the experiments, no neurological and psychiatric disorders, no ongoing diets, no regular intake of medication, no smokers. Additionally, participants were asked to abstain from ingesting any food or caloric drink within the 3 hours prior to the experiments. In order to assess normal tasting and smelling abilities, participants passed two clinical tests: ‘Taste Stripes’ and ‘Sniffin’ Sticks’ (Burghart Messtechnik, Wedel, Germany). All participants gave written informed consent according to the local ethics committee of Otto-von-Guericke-University Magdeburg.

### Stimuli

We used four distinct tastants (bitter, salty, sour, sweet) plus a neutral solution (artificial saliva). Solutions were provided by the central pharmacy of the medical faculty of Otto-von-Guericke-University Magdeburg. Stimuli were obtained from the following basis solutions (= 100%): 600 mM NaCl (salty), 1 M glucose (sweet), 0.1 mM quinine hydrochloride (bitter) and 8 mM citric acid (sour). The neutral solution consisted of 5 mM KCl and 0.5 mM NaHCO_3_, the same chemical compounds as saliva (O’Doherty et al., 2001) though at lower concentration as O’Doherty et al.(2001) based on own piloting. With the exception of the neutral taste, all stimuli were presented at two different concentrations (low and high), which were chosen via pilot testing in order to guarantee a discernible difference between the two concentrations. The resulting stimuli were obtained according to the following percentages of basis solution dissolved with pure water (low/high): 10 %/70 % salty, 20 %/80 % sweet, 20 %/100 % bitter, 10 %/80 % sour.

Stimuli were delivered through an automated stimulation device: Gustometer GU002 (Burghart Messtechnik, Wedel, Germany). The Gustometer allows the control of stimulus duration and concentration and massively reduces undesirable somatosensory stimulation given that stimuli are sprayed via compressed air directly onto the tip of the participants’ tongue. The Gustometer delivers the stimulus at ca. 40°C resulting in approximately the body temperature of the stimulus upon tongue contact. The device was controlled through Matlab 2012b (Mathworks, Inc., Natrick, MA) via custom made scripts and Psychophysics Toolbox (Brainard, 1997) running on a Windows 7 environment.

Stimuli were delivered to the participant through a pump hold by a custom made plexiglass scaffolding mounted above the head coil. Participants were asked to protrude the tip of their tongue and enclose it with their lips in order to prevent to ingest the liquid. None of the participants reported any physical discomfort related to this procedure. The liquid was then absorbed by a tissue placed below the participant’s mouth. This procedure was adopted to avoid any motion and swallowing related artifacts and most importantly to isolate taste-related activity; while avoiding or holding constant other confounding factors (e.g. temperature, stimulation of oral cavity etc.).

### Experimental Procedure

Participants took part in two scanning sessions with a minimum time interval of 1 day and a maximum of 7 months. Before and after any scanning session, participants – while laying in the scanner – were asked to identify each one of the stimuli presented during the experiment, resulting in four distinct identifications per taste (see supporting Fig. S1a-b for descriptive results).

The main experiment consisted of four runs, each composed of 45 trials. Each run consisted of sequences of stimuli comprised of all tastes in both concentrations plus the neutral taste. All sequences had to fulfill three basic constraints: 1) each stimulus was presented 5 times within each run, 2) the same stimulus could never be presented twice in a row, 3) each combination of two consecutive stimuli was presented at least once during the entire experiment by using De Brujin sequences.

During scanning, each trial started by concurrently displaying the word “Schmecken” (“Taste” in German) and spraying the tastant on the tongue of the participant. The stimulation consisted of four distinct sprays (250 ms each) interspersed with four pauses (250 ms each) for a total duration of 2 seconds; each spray released 50 μl of tastant, summing up to a total of 200 μl per trial. The tastant delivery was followed by a rating scale which assessed intensity or pleasantness for the first and second experiment, respectively (“Wie intensiv/angenehm empfinden Sie diesen Geschmack?”, “How intense/pleasant does this taste?” in German). Participants were asked to provide their rating on a nine-point-scale ranging from 1: “No perception” to 9: “Extremely intense”, or “Extremely unpleasant” to “Extremely pleasant” displayed on the screen. In order to respond, participants slid a green cursor through the numbers of the scale by pressing two buttons with the index and middle finger of the right hand. To confirm their choice, participants pressed a third button with the thumb. For each rating scale, the green cursor was randomly located on a different digit upon scale display to avoid biasing. In case a participant was unable to provide an answer within 6 seconds after the appearance of the scale, the response was considered invalid. Immediately after the rating scale, the word “Rinse” was presented on the screen together with 2 seconds of sprayed water to rinse the tongue from the previous tastant. The following trial started after an intertrial interval of 2 seconds (total trial duration 12 s).

### Neuroimaging Data Collection

Scanning was conducted on a Siemens Prisma 3 Tesla system with a 32-channel head coil for signal reception. T1-weighted structural images were acquired with an MPRAGE sequence using the following parameters: 1 x 1 x 1 mm^3^ voxel size, 256 x 256 x 192 matrix, 2.82 ms echo time, 2.5 s repetition time, 1.1 s inversion time, 7° flip angle, 140 Hz/pixel bandwidth, 7/8 partial Fourier, parallel imaging with a GRAPPA factor of 2, and 5:18 min scan duration.

For functional imaging we opted for a combination of reduced field-of-view (rFOV) gradient echo EPI and parallel imaging in order to minimize signal dropout and geometric distortion. This technique is part of the Siemens “Advanced fMRI” work-in-progress software. During each of the 4 functional runs we acquired 280 volumes using the following parameters: 24 slices, 1.8 mm slice thickness, 0.9 mm interslice gap, ascending slice excitation order, 135 x 240 mm^2^ rFOV, 1.25 x 1.25 mm^2^ in-plane voxel size, 30 ms echo time, 2 s repetition time, 90° flip angle, 1 mT/m·ms z-shim, 0.73 ms echo spacing, and GRAPPA factor of 2. The transversal slice block was tilted 20° with respect to the AC-PC line.

To facilitate the registration and normalization of the rFOV we additionally acquired 10 whole brain EPI volumes. In order to keep geometrical distortions within the rFOV part of the whole brain data set identical to the functional scans the shim settings were copied and the FOV and matrix size in phase direction as well as the GRAPPA factor were doubled. A Siemens auto-align-algorithm was used to automatically select the identical slices during the second scanning session.

## Data Analysis

### Behavioral data

Ratings of intensity and pleasantness were analyzed with the same statistical procedure: We first computed the median across all the ratings separately for each experimental condition. Each dataset was then subjected to a repeated-measures ANOVA with factors: *Taste category* (bitter, salty, sour, sweet and neutral) and *Intensity* (high and low). Post-hoc t-tests were subsequently performed to statistically test potential differences among taste ratings. In order to correct for multiple comparisons, Bonferroni correction was applied.

Brain-behavior correlations were computed using Pearson’s correlation coefficients and Bonferroni corrections. Effects of *taste category* and *intensity* were analyzed using repeated measures ANOVAs with the factors *hemisphere* and *taste category*, separately for low and high concentrations. All analyses were performed by using the R package *stats* v3.6.0 within the R environment (v3.4.4).

### fMRI data pre-processing

fMRI analysis was conducted within a computational cluster in a Debian environment; analysis tools were obtained through NeuroDebian (Halchenko and Hanke, 2012). Both datasets were subject to the identical pre-processing procedure: Prior to any preprocessing, 4 dummy scans, collected at the beginning of each scanning session were removed from each dataset. Motion correction was performed by re-aligning, each volume of both experiments to the first volume of experiment 1 via a rigid body transformation (MCFLIRT, FSL 5.09).

For noise reduction, time series were temporally filtered by means of a band pass filter (cutoffs: 4 Hz, 150 Hz). Additionally, each volume was spatially smoothed by applying a FWHM Gaussian kernel of 4 mm via Nilearn 0.5.0 (Abraham et al., 2014).

### Multivariate Pattern Analysis (MVPA)

Decoding analysis was performed with PyMVPA (Hanke et al., 2009) and custom Python scripts. Prior to the MVPA, each time series was fitted on a voxel by voxel basis with a General Linear Model (GLM) using Nipy 0.4.2 (Millman and Brett, 2007).

The GLM was composed of five “taste” regressors (neutral, bitter, salty, sour, sweet; trial onset aligned with start of taste delivery, duration 2 s) convolved with a canonical hemodynamic response function (HRF) with temporal derivatives. Motion estimates (6 DOF) were included as additional nuisance regressors. In order to subtract potential artifacts common to each taste condition, GLM parameters for each taste were contrasted runwise against the neutral condition and the resulting t-contrasts were used as features for classification.

We then applied a Sparse Multinomial Logistic Regression classifier (SMLR, Krishnapuram et al., 2005) and a leave-one-out cross-validation procedure. The type of folding used for cross-validation changed according to the aim of the specific classification performed. For the cross-experiment classification, the SMLR was, in turn, trained on one experiment and tested on the other one; for the cross-concentration classification, the SLMR was trained on low concentration data and tested on the high concentration data and vice versa. For the within experiment classifications (separate classifications for high and low concentrations for both experiments), the SMLR was, in turn, trained on three out four runs and tested it on the remaining run. Classification was restricted to voxels in the insula using the brain parcellation provided by Fan et al., 2016, with a 50% cut-off criterion for the probability maps. Insular ROIs included the following left and right subregions: dorsal dysgranular insular (dId), dorsal agranular insular (dla), ventral agranular insular (vla), dorsal granular insular (dlg), ventral granular insular (vlg/vId) and hypergranular insular (G) cortex.

Decoding was performed through a Searchlight approach (Kriegeskorte et al., 2006) with a sphere’s radius of 4 voxels. The primary outcome of the classification analysis were the taste-category-wise probability estimates of the SMLR classifier, for each searchlight sphere and each data fold.

### MVPA statistical analysis

In order to assess the degree of evidence for a response to a specific presented taste, we computed the following ratio for each taste and searchlight sphere:

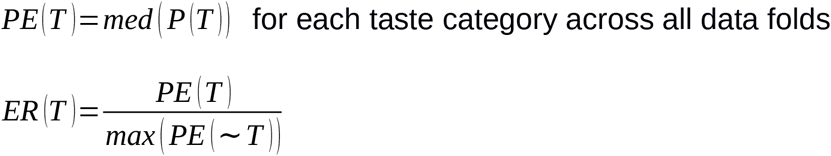

Where: PE(T) denotes the median of the SMLR probability estimates P(T) for one specific taste across all trials in which this particular taste was presented. PE(~T) is the median of probability estimates for the same taste based across all trials in which this particular taste was not presented (i.e. a different taste was presented). The rationale adopted for the Evidence Ratio (ER) resembles the concept of the Bayes factor, by comparing the alternative PE(T) and null PE(~T) hypothesis to assess the goodness of the model of interest.

ER computation yields a single map for each taste. In total, four evidence ratios, one for each taste, were calculated per data fold. The obtained maps were then statistically evaluated at the group level following a bootstrapped permutation analysis proposed by Stelzer et al. (2013), implemented in PyMVPA, and briefly summarized here: In order to assess the null distribution of results, we computed 50 additional “chance” result maps for each participant, using the identical analysis setup and data folding strategy, while permuting the taste labels within the training data. For group-level inference, all individual maps were spatially transformed and re-sliced, via FSL FLIRT, from native image space into MNI space with a 2 mm isotropic voxel size.

The empirical group average map was thresholded using a variable voxel-wise cut-off, corresponding to p < 0.001 of the distribution of results from 10000 bootstrap samples of group average maps, computed from randomly drawn “chance” maps, one for each participant. Likewise, group-level cluster size was statistically tested (p < 0.05) based on the distribution of cluster sizes across the 10000 bootstrap samples, after applying the same p < 0.001 voxel-wise threshold to all “chance” group average maps.

### Quantifying tuning maps

In order to quantify the degree of selectivity for each taste or a combination of taste categories, we computed an index of preferential tuning for each sphere. Tuning indices (TI) were based on ERs and calculated by using the single taste ERs (e.g. salty) or by combining the ERs of different tastes (e.g. bitter and salty). Importantly, TIs were calculated both on the group-level and single subjects’ statistical maps according to the following equation:

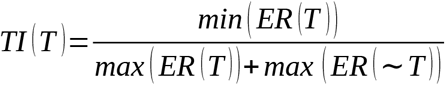

Where: ER(T) indicates the evidence ratio for – a single taste or a combination of tastes of interest – and ER(~T) indicates the evidence ratios of the remaining tastes. In case of single taste tuning, min(ER(T)) and max(ER(T)) are simply ER(T). The choice of taking the minimum of ER(T) – in the numerator – and maximum of ER(T) – in denominator – becomes relevant only when we want to evaluate the selectivity of a combination of tastes (e.g. bitter **and** salty). Selecting the smaller ER(T) in the numerator and the bigger ER(T) in the denominator prevents that two highly different ER values would produce a TI above 0.5, which would indeed provide a misleading characterization of that sphere’s preference. Tuning indices guarantee that a given sphere can be considered as preferentially tuned to a taste or a combination of tastes only if the tuning index is equal or bigger than 0.5. Hence, they provide a rather conservative measure (range 0-1) either in favor – values above or equal to 0.5 – or not in favor of the selected taste – values below 0.5. Additionally, to avoid selecting extremely low but significant evidence ratios, we selected spheres with ER(T) equal or larger than 3 for at least one taste. This threshold was motivated by the analogy with the Bayes factor, where values equal or larger than 3 are considered indicative of a moderate evidence (Kass and Raftery, 1995).

### Bland and Altman plots

Bland and Altman plots (Bland and Altman, 1999, see Figure 4) represent the amount of agreement between high and low concentration evidence ratios in the cross experiment classification. In order to construct such plots, we first assembled the ERs of a specific taste contained in the ROI of all subjects. Given the high variability of the ER values, we log-transformed (log2) the data to merely improve results visualization. The x axis of the plots represents the average between high and low concentration ERs. The y axis represents the difference between high and low concentration ERs. The blue dots are simply the result of the aforementioned computations, thus representing a general agreement between high and low concentration ERs. The red dots represent, instead, the agreement between co-active spheres in high and low concentration ERs that exhibited also a tuning index equal or larger than 0.5. Note that there is no circularity in this approach given that ERs with tuning indices equal or larger than 0.5 can have completely different values and are not conditioned by any common constraint, except for the requirement to have an ER value equal or larger than 3 (see the Quantifying tuning maps section). Hence, the clustering around 0 along the y-axis provides critical additional information about these coactive spheres.

## Acknowledgements

EP, MH and TN were funded by DFG-SFB779/TPA15.

## Author contributions statement

KMB established the experimental methods and acquired the data. EP and MH analyzed the data. TN and MH designed the experiments. All authors wrote the manuscript.

## Data/Code availability

The datasets/code supporting the current study have not yet been deposited in a public repository because of planned additional publications but are available from the corresponding author on request.

