## supporting material for "Information-based taste maps in insular cortex are shaped by stimulus concentration"

Supporting Material contains

2 Supporting Tables S1, S2; 3 Supporting Figures S1-S3

**Supporting Table S1.**

| **Pleasantness** | | | | |
| --- | --- | --- | --- | --- |
| **High versus high** | | | **Low versus low** | |
| **Tastes** | ***t* - value (df=23)** | ***p* val*ue***  **(Bonferroni corrected)** | ***t* - value (def=23)** | ***p* value**  **(Bonferroni corrected)** |
| Bitter - Salty | -0.1618515 | 1 | 1.631 | .51 |
| Bitter - Sour | -1.722031 | .51 | -0.81626 | 1 |
| Bitter - Sweet | -9.909251 | <.001 | -4.22067 | .01 |
| Bitter - Neutral | -8.29141 | <.001 | -3.49772 | .02 |
| Salty - Sour | -1.802176 | .51 | -3.112504 | .05 |
| Salty - Sweet | -8.2239 | <.001 | -5.740817 | <.001 |
| Salty - Neutral | -3.947769 | .01 | -3.866523 | .01 |
| Sour - Sweet | -6.696908 | <.001 | -2.30754 | .24 |
| Sour - Neutral | -2.302078 | .24 | -0.6925073 | 1 |
| Sweet - Neutral | 6.486383 | <.001 | 3.148802 | .05 |
| **High versus Low** | | | | |
| **Tastes** | ***t*-value (df=23)** | | ***P*-value** | |
| Bitter - Bitter | -9.525788 | | <.001 | |
| Salty - Salty | -2.506884 | | .18 | |
| Sour - Sour | -4.089396 | | .01 | |
| Sweet - Sweet | 5.910092 | | <.001 | |

| **Intensity** | | | | |
| --- | --- | --- | --- | --- |
| **High versus high** | | | **Low versus low** | |
| **Tastes** | ***t*–value (df=23)** | ***p* value** | ***t–*valu*e* (df=23)** | **p value** |
| Bitter - Salty | -5.800154 | <.001 | -4.495521 | <.001 |
| Bitter - Sour | -3.185445 | .02 | -5.065961 | <.001 |
| Bitter - Sweet | -2.46021 | .09 | -1.098434 | .28 |
| Bitter - Neutral | 7.440002 | <.001 | 3.301131 | .02 |
| Salty - Sour | 5.385858 | <.001 | 2.118493 | .14 |
| Salty - Sweet | 6.911529 | <.001 | 3.750918 | .01 |
| Salty - Neutral | 14.84924 | <.001 | 6.351713 | <.001 |
| Sour - Sweet | 1.762983 | .18 | 2.984682 | .03 |
| Sour - Neutral | 11.02563 | <.001 | 6.94982 | <.001 |
| Sweet - Neutral | 11.23475 | <.001 | 3.768823 | .01 |
| **High versus Low** | | | | |
| **Tastes** | ***t*-value (df=23)** | | ***p*-value**  **(Bonferroni corrected)** | |
| Bitter - Bitter | 8.494864 | | <.001 | |
| Salty - Salty | 10.67065 | | <.001 | |
| Sour - Sour | 9.690101 | | <.001 | |
| Sweet - Sweet | 12.62226 | | <.001 | |

**Supporting Table S1. Pairwise comparisons for pleasantness and intensity judgments:** The upper table reports pairwise t-tests comparisons for pleasantness ratings at different intensities (see main text for results of repeated ANOVA). Results indicate that high concentration and low concentration bitter and salty stimuli were perceived as less pleasant than neutral and sweet. Sweet was perceived as more pleasant than neutral. Moreover highest changes in valence due to concentration were found for sweet and bitter. Lower table reports pairwise t-tests comparisons for intensity ratings. Note that all low- and high-concentration tastes were significantly different from neutral and that all high-concentration tastes were judged as more intense than their low-concentration counterparts. All p-values were Bonferroni corrected. See Supporting Fig. S1c-d for condition-wise group means.

**Supporting Table S2.**

| **A. High Concentration**  **Double combinations** | **Non-responders** | | | |
| --- | --- | --- | --- | --- |
| Taste combinations | Median (%) | SE | Left | Right |
| Bitter salt | 0.47 | 0.27 | 2 | 2 |
| Bitter sour | 0.37 | 0.28 | 3 | 3 |
| Bitter sweet | 0.46 | 0.26 | 1 | 2 |
| Salt sour | 0.41 | 0.25 | 2 | 2 |
| Salt sweet | 0.51 | 0.22 | 2 | 2 |
| Sour sweet | 0.45 | 0.21 | 2 | 2 |

| **B. High concentration**  **Triple combinations** | **Non-responders** | |
| --- | --- | --- |
| Taste Combinations | Left | Right |
| Bitter salt sour | 13 | 19 |
| Bitter salt sweet | 11 | 13 |
| Bitter sour sweet | 15 | 13 |
| Salt sour sweet | 13 | 15 |

| **C. Low concentration**  **Double combinations** | **Non-responders** | | | |
| --- | --- | --- | --- | --- |
| Taste combinations | Median (%) | SE | Left | Right |
| Bitter salt | 0.25 | 0.18 | 2 | 5 |
| Bitter sour | 0.19 | 0.15 | 5 | 6 |
| Bitter sweet | 0.46 | 0.37 | 1 | 1 |
| Salt sour | 0.18 | 0.21 | 3 | 7 |
| Salt sweet | 0.30 | 0.17 | 2 | 2 |
| Sour sweet | 0.37 | 0.19 | 5 | 4 |

| **D. Low concentration**  **Triple combinations** | **Non-responders** | |
| --- | --- | --- |
| Taste Combinations | Left | Right |
| Bitter salt sou | 18 | 22 |
| Bitter salt sweet | 17 | 19 |
| Bitter sour sweet | 17 | 18 |
| Salt sour sweet | 17 | 16 |

**Supporting Table S2:** Descriptive Statistics for subject-specific double- and triple taste-sensitive spheres indicated in percent of all insular spheres (median +/- SE). Non-responders denote the number of participants with no significant clusters in left and right insula. For triple combinations, no median were calculated as more than half of subjects showed no significant responses (i.e. median would have been 0).

**Supporting Figure S1.**

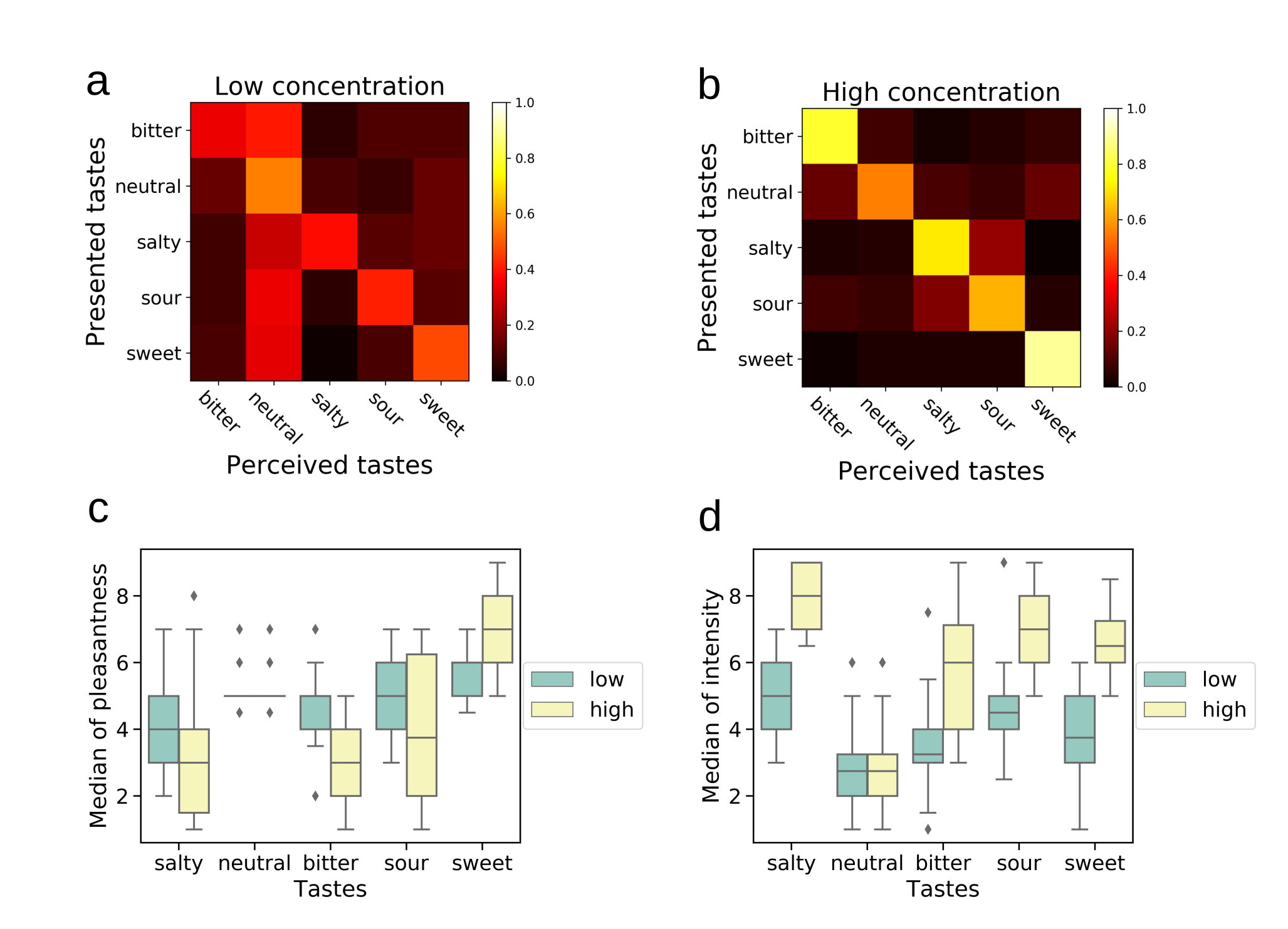

**Supporting Figure S1. Identity judgements**. (a-b) Confusion matrices depict response ratios for perceived tastes (x-axis) relative to presented tastes (y-axis). Left heat map depicts low concentration tastes; right heat map depicts high concentration tastes. Diagonal represents correct identifications, which always showed the highest response frequencies relative to all other response options (accuracies at 37 percent or higher) with the exception of low-concentration bitter. Note that the neutral taste (2^th^ row) was identical for both maps, as neutral was by definition presented without intensity manipulation. (c-d) Valence and intensity judgements inside scanner**.** The statistical comparisons of the presented data can be found in Supporting Table 1 for statistical comparisons. (c) Boxplot depicts condition-wise median valence ratings across participants. Boxes represent interquartile ranges (Blue boxes= low concentration; yellow box=high concentration). Dots represent outliers. See Supporting Table 1 for statistical comparisons. (d) Boxplot represents median intensity ratings across participants. See Supporting Table 1 for statistical comparisons. Neutral was presented only at one concentration and duplicated in the box plots for display purposes.

**Supporting Figure S2.**

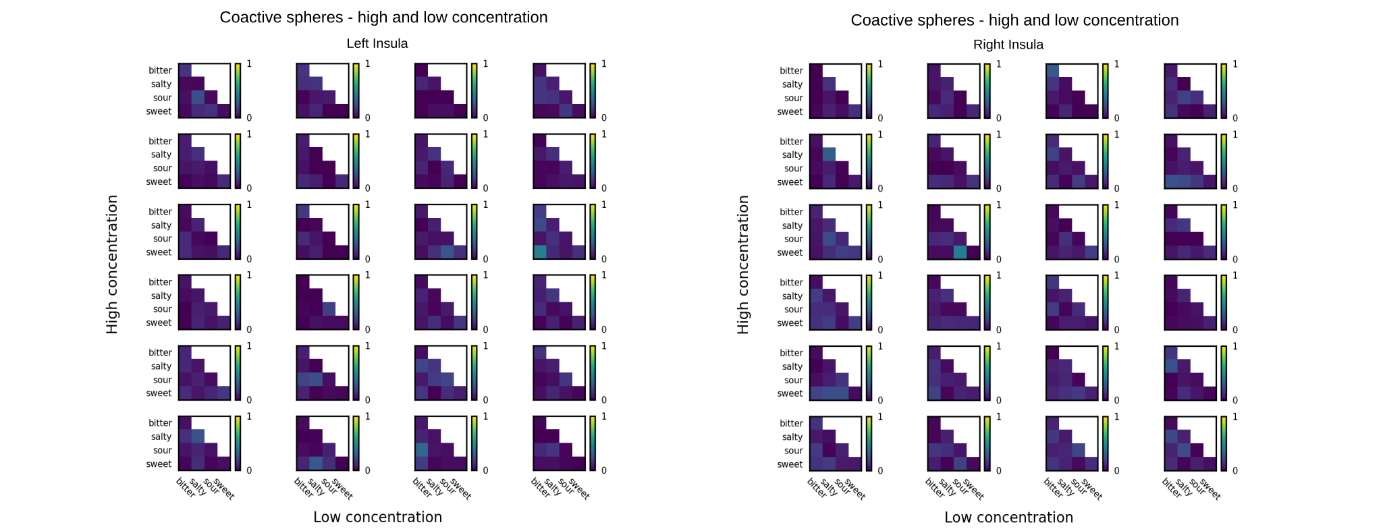

**Supporting Figure S2. Coactive spheres across concentrations in left and right insula.** Heat maps depict, for each of the 24 subjects, the index of narrowly tuned spheres coactive in low and high concentrations across experiments. Upper left heat map represents overlap in subject 1, followed by subject 2 to the right. Coactive spheres are expressed as an index of overlap, where a 0 indicates the absence of overlap and 1 complete overlap. The main diagonal represents spheres coding for the identical taste category across concentrations. Values outside the main diagonal indicate a switch in taste preference as a function of concentration.

**Supporting Figure S3.**

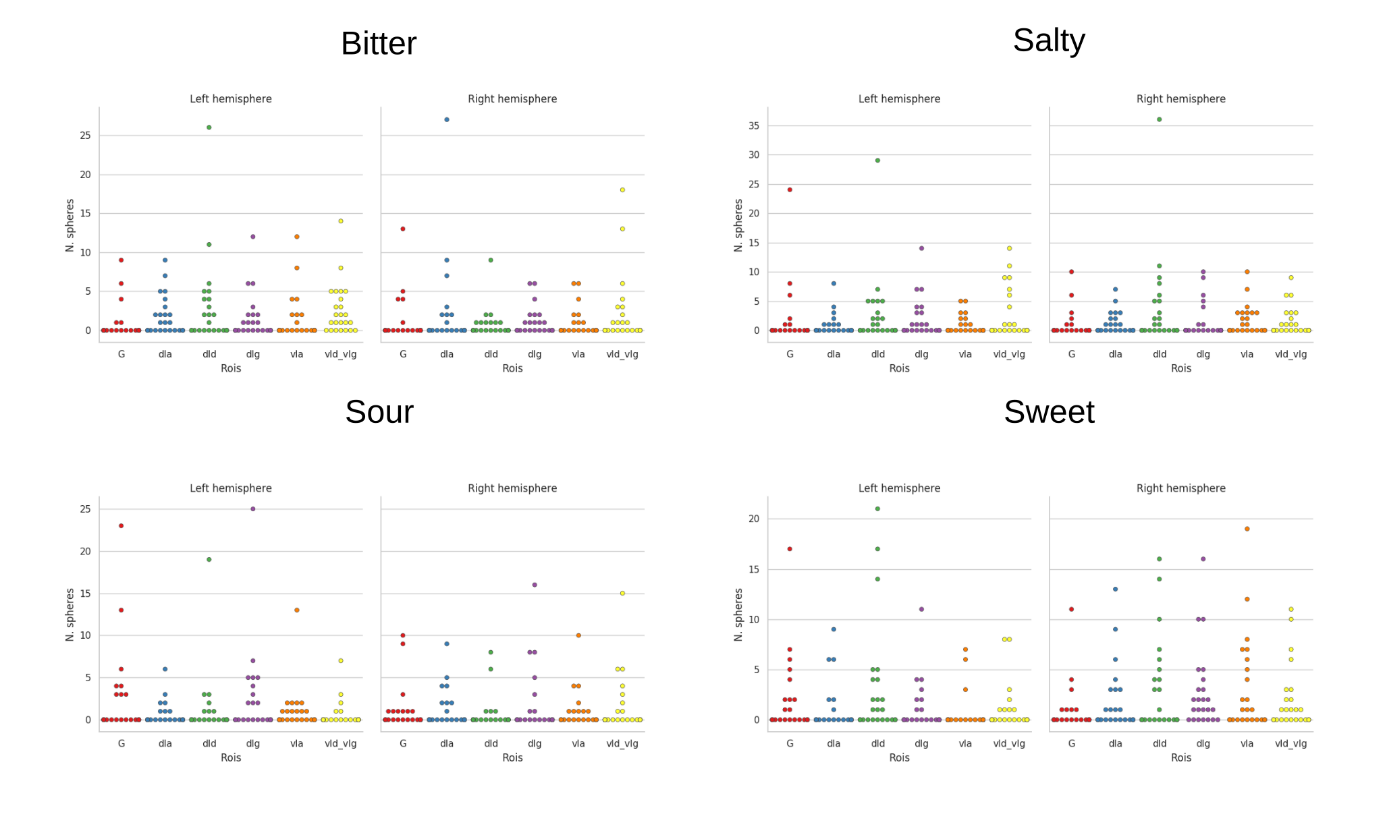

**Supporting Figure S3. Distribution of concentration-invariant narrowly tuned spheres across insular subregions:** Swarm-plots depict the number of spheres (y-axis) per each insular subregion tuned to a specific taste. Dots represent single subjects. The sphere count was obtained from the single subject tuning maps of each specific taste. Note that subjects could contribute spheres to more than one subfield. Insular subfields: Dorsal dysgranular insular (dId), dorsal granular insular (dlg), dorsal agranular insular (dla), ventral agranular insular (vla), ventral granular insular (vld/vIg) and hypergranular insular (G) cortex (Fan et al., 2016).
